# Cell proliferation depends on the direct binding between PKM2 and AKAP-Lbc

**DOI:** 10.1101/103333

**Authors:** Xinping Chen, Graeme K. Carnegie

## Abstract

The M2 form of the glycolytic enzyme pyruvate kinase (PKM2) has generated much interest recently due to its important role in tumor metabolism. A yeast two-hybrid screen carried out by the Alliance for Cell Signaling suggests that PKM2 interacts with A-Kinase Anchoring Protein (AKAP)-Lbc.

AKAP-Lbc (also known as AKAP13) is a scaffold protein that integrates signaling through multiple enzymes including protein kinases A and D and the small G protein Rho. AKAP-Lbc was originally identified in leukemic blast cells, and multiple reports implicate AKAP-Lbc in breast, prostate and thyroid cancers, however the role of AKAP-Lbc in cancer biology is not understood.

Co-immunoprecipitation, pulldown and Bimolecular Fluorescence Complementation (BiFC) data indicate that PKM2 interacts with AKAP-Lbc. Mapping experiments indicate that PKM2 directly interacts with amino acid residues 1923-2817 of AKAP-Lbc. By disrupting the interaction between the two proteins with the expression of the AKAP-Lbc fragments, our data suggest that the binding between PKM2 and PKA plays a critical role in cell proliferation. The work indicates that the binding between AKAP-Lbc and PKM2 may be an important target to treat some cancers by reducing the cell proliferation.

## Introduction

Tumor cells prefer glycolysis instead of mitochondrial oxidative phosphorylation to generate ATP,, the Warburg effect (Vander Heiden, Cantley et al. 2009). As a crucial enzyme, PKM2 produces ATP in glycolysis, and PKM2 has been reported to be important for the proliferation and growth and resistance to some drug treatment of cancers (Shi, Li et al. 2010, Wong, De Melo et al. 2013, Rahman and Hasan 2015), and the correlation of PKM2 to oncogenesis has been reported in multiple cancers, such as colorectal cancer (Dayton, Jacks et al. 2016, Dong, Mao et al. 2016)., It is thus worthwhile to understand how PKM2 is distributed and regulated.

Based on the yeast two-hybrid screen by the Alliance for Cell Signaling, we proposed that PKM2 may interact with AKAP-Lbc. AKAP-Lbc has been implicated to play a role in some cancers (ref expected here), but the details of this role remain to be specified.

To confirm the potential interaction between these two proteins, we selected molecular and biochemical methods to explore the underlying relations of these two proteins.

## Methods

### Cell culture and transfection

Cells were cultured in DEME added with 10% of FBS as reported (Wang and Carnegie 2013). For transfection, each well in a 6 well plate, received 133 μL of milliQ water containing 2-20 μg of DNA, and 17 μL of 2.5 M CaCl2.

### Cell lysis

HEK293T cells grown in a 6-well plate received 100 μL per well, while cells grown in a 10 cm petri dish received 500 μL of IP/lysis buffer (10 mM sodium phosphate buffer (pH 6.95), 150 mM NaCl, 5 mM EDTA, 5 mM EGTA, and 1% Triton X-100). Plates were put on ice for several minutes and rocked detaching cells from the plates. Cells were then centrifuged to remove the debris (Burmeister, Wang et al. 2015).

### Antibodies

PKM2 Antibody was from Cell Signaling Technology (#3198), anti-Flag monoclonal antibody was from Sigma (F1804), and anti-HA monoclonal one was purchased from COVANC (16B12).

### Confocal imaging

Imaging was done with a Carl Zeiss LSM 510 mounted on an Axiovert 100 M microscope (Burmeister, Wang et al. 2015).

### Expression of protein in BL21 or DH-5alpha cells

25 ml bacterial cultures were grown in LB medium overnight at 37°C, and diluted to OD600 between 0.4–0.6. Some of the culture was reserved as a negative control for protein expression. GST-fusion or His-tagged protein was induced with IPTG at the final concentration as 50 μM for 3 h at 32°C. Bacteria were spun down at 6,000 g at 4°C for 15 min. Medium was decanted and 10-20 ml IP/lysis buffer or with *E. coli* lysis buffer (50 mM NaH_2_PO_4_, 300 mM NaCl, 10 mM imidazole, pH 8.0) with added protease inhibitors. Pellets were sonicated (Sonifier 250, Branson: Timer-Hold; Duty cycle-constant, output control-2.5, 10 s each for twice) until no pellet remained. Lysates were transferred to small plastic tubes and centrifuged at 20, 000g at 4 °C for 30 min. Supernatants were transferred to 15 ml Falcon tubes 130 μL Ni-NTA (Lot: 139281394, QIAGEN) or GST bead (Glutathione sepharose^TM^ 4B Lot: 10039158, GE) slurry was added. After overnight incubation,. tubes were centrifuged for 30s at 3,000g. Beads were transferred to Eppendorf tubes with lysis buffer five times (for GST beads, using the lysis/IP buffer without EDTA and EGTA; for Ni-beads, wash buffer contained 50mM NaH2PO4,300mM NaCl, 20mM imidazole, pH 8.0). 20 μL of the GST bead slurry was boiled with loading buffer and run on a 10% SDS-page gel. For Ni-NTA beads, proteins were eluted with elution buffer (50 mM NaH2PO4,300 mM NaCl, 250 mM imidazole, pH 8.0) three times, and and 20 µL run on the gel. Lanes with 1, 5 and 10 μg BSA were run as control. Proteins were to PVDF membranes and stained with Ponceau S (Lot 091M4378V, Sigma) solution.

## Cell proliferation analysis

5 × 104 cells were seeded in triplicate in 6-well plates, and accurate cell counts were obtained every 24 h using a Coulter particle analyzer for a 3–5-day period. Time zero was taken 16 h after seeding. The standard MTT assay was applied according to the instruction protocol (Sigma).

## Results

### Presence of PKM2 and AKAP-Lbc in cancer cells

To explore whether PKM2 and AKAP-Lbc interact in cancer biology, it is important to confirm that both proteins are expressed in cancer cells. We chose several cancer cell lines to examine the possibility. Firstly, we tested several commonly used cell lines to detect the expression of both proteins. As shown in Figure 1A, we observed co-expression in three cell lines, including A549 (Lung cancer), HeLa (cervical cancer) and 293T cells (note that 293T cells are not considered true cancer cells (Graham, Smiley et al. 1977)) and the 293T cellswere applied to explore the interaction in most of the current work. Further, the two proteins were co-expressed in other cancer cell lines, NLF (Neuroblastoma), Lan1 (Neuroblastoma), NB69 (Neuroblastoma), IMR-5 (Neuroblastoma), SK-N-AS (Neuroblastoma), SY5Y (Neuroblastoma), MIA PaCa (pancreatic carcinoma) and CHP-134 (Neuroblastoma) (Figure 1B), suggesting that the two proteins are present in multiple cancers.

**Figure 1.**
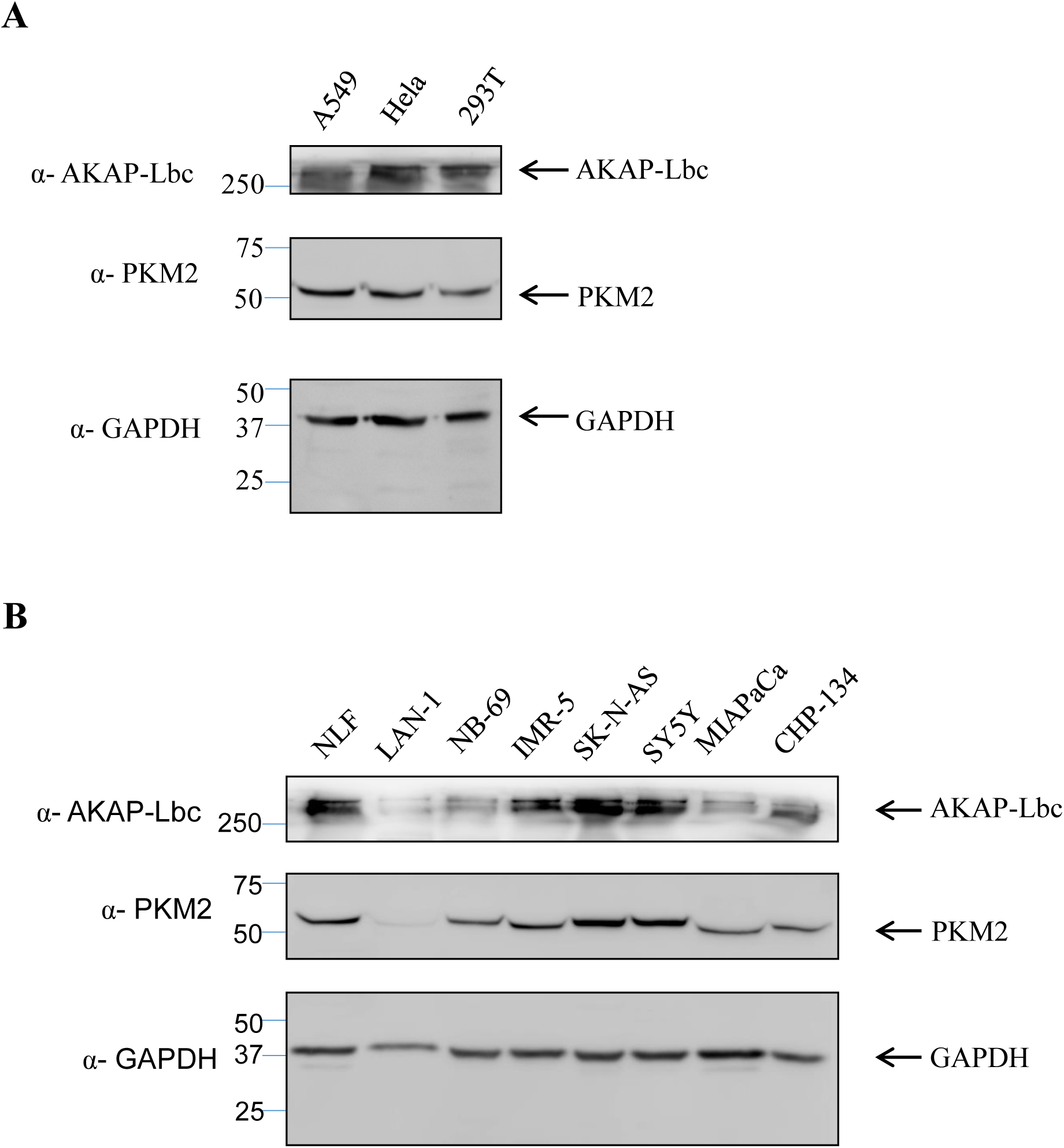
AKAP-Lbc and PKM2 expression in Hek293T, Hela and A549 lung carcinoma and neuroblastoma cell lines. AKAP-Lbc and PKM2 expression in A) Hek293T, Hela and A549 lung carcinoma cells, as well as in B) eight neuroblastoma cell lines.

### The interaction of PKM2 and AKAP-Lbc in HeLa cells

Since we found that the two proteins are expressed in multiple cancer cell lines, it is intriguing to investigate if they interact in cancer cells. To address this question Flag-AKAP-Lbc was expressed in HeLa cells, and cell extract was incubated with anti-Flag beads to pull down Flag-AKAP-Lbc and potential binding partners. We found that Flag beads pull down both Flag-AKAP-Lbc and endogenous PKM2 in the cell extract (Figure 2A). Though there is still some signal in the non-transfection control for PKM2, the low level of PKM2 is considered non-specific binding to the beads. To further examine if endogenous AKAP-Lbc interacts with PKM2 in HeLa cells, we used anti-AKAP-Lbc antibody to attempt co-immunoprecipitation (co-IP). Consistent with the anti-Flag IP result, we observed that anti-AKAP-Lbc IP pulled down both AKAP-Lbc and PKM2, while control IgG beads did not, indicating that the two proteins interact in cancer cells.

**Figure 2.**
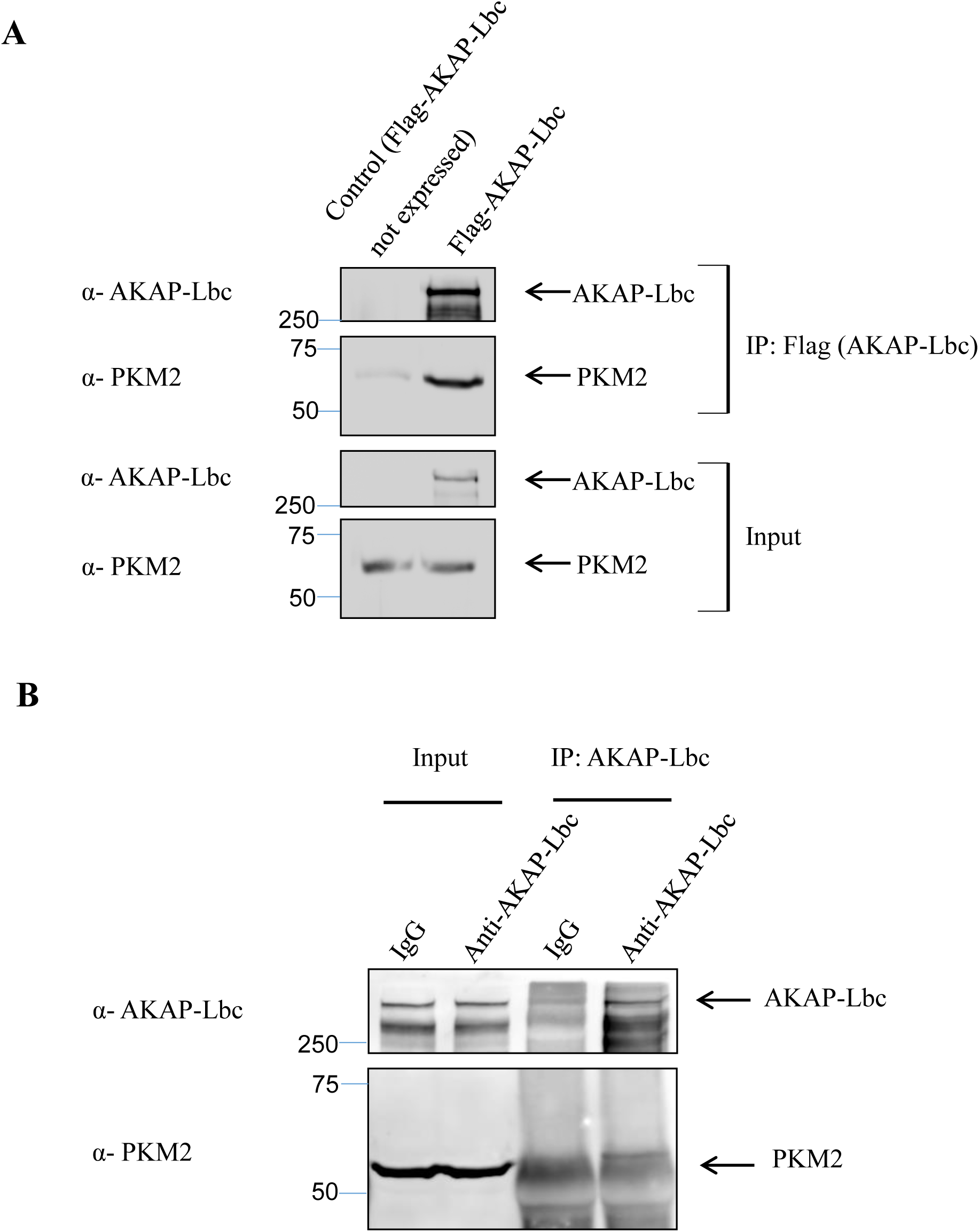
PKM2 binds to AKAP-Lbc in Hela calls. A) Flag-AKAP-Lbc wasexpressed in HeLa cells and immunoprecipitated. Western blot demonstrates co-immunoprecipated of PKM2 with Flag-AKAP-Lbc. B) Co-immunoprecipitation of endogenous PKM2 with endogenous AKAP-Lbc in HeLa cells.

### Visualization of the interaction of PKM2 and AKAP-Lbc in cells

To visualize AKAP-Lbc/PKM2 in live cells, we used Bimolecular Fluorescence Complementation (BiFC). Figure 3A schematically shows the BiFC stratedgy: protein-protein interaction brings two separated Venus fragments tagging the two proteins together, reconstituting Venus fluorescence. To validate the system, we set several negative controls that should not show fluorescence, including single expression of AKAP-Lbc, PKM1 and PKM2, or the combined expression of two genes, such as HRas and PKM1 (Figure 2B). Compared to the negative controls, we observed strong fluorescent signal in the cells co-expressing VN-AKAP-Lbc and VCC-PKM2 while weak one in the cells expressing VN-AKAP-Lbc and VCC-PKM1, suggesting the interaction between AKAP-Lbc and PKM2 is stronger than that of AKAP-Lbc and PKM1. Consistent with co-immunoprecipitation and bead pull-down, BiFC results support interaction between AKAP-Lbc and PKM2 in live cells, prompting us to ask how they interact with each other.

**Figure 3.**
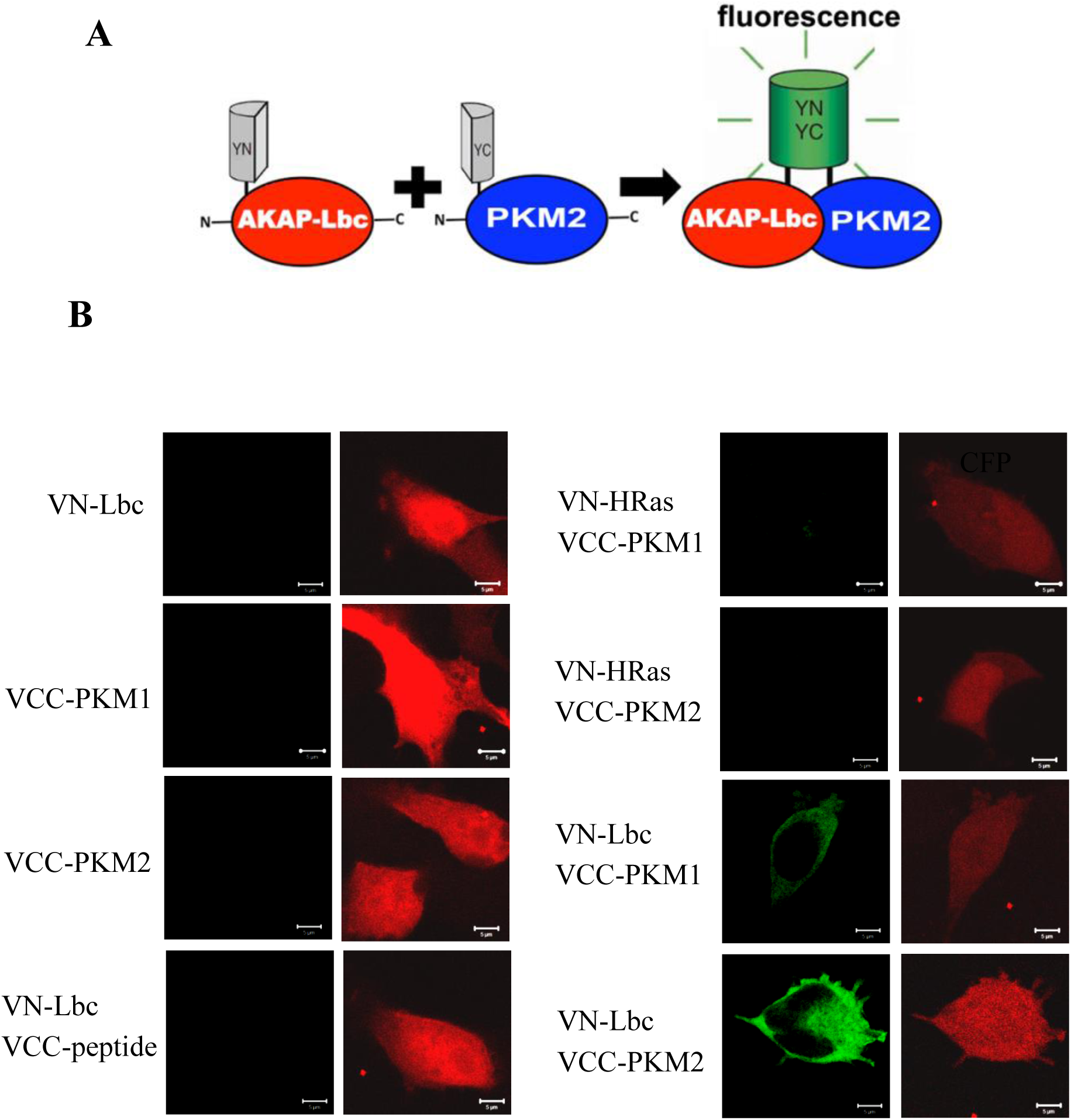
Visualization of AKAP-Lbc-PKM2/PKM1 interaction inside cells by Bimolecular Fluorescence Complementation (BiFC). Schematic diagram illustrating the BiFC assay. The venus (green) fluorescent protein is split into two non-fluorescent halves which are fused to AKAP-Lbc and PKM2/PKM1. Specific protein-protein interaction results in a functional venus (green) fluorescent protein. **B)** HEK 293 cells were co-transfected with VN-AKAP-Lbc and VC-PKM2/PKM1 and CFP (pseudo-colored red), as a marker for transfected cells. Cells were fixed 24 hrs after transfection and imaged. No BiFC (green) fluorescence is observed in cells expressing either VN-AKAP-Lbc or VC-PKM2/PKM1 alone, or when VN-AKAP-Lbc is co-expressed with a non-interacting control VC-peptide. Similarly, no BiFC (green) fluorescence is observed when VC-PKM2/PKM1 is co-expressed with a control VN-protein (VN-HRas), whereas specific interaction of VC-PKM2/PKM1 with VN-AKAP-Lbc results in fluorescence. Equal protein expression was determined by Western blot (data not shown).

**Figure 3.**
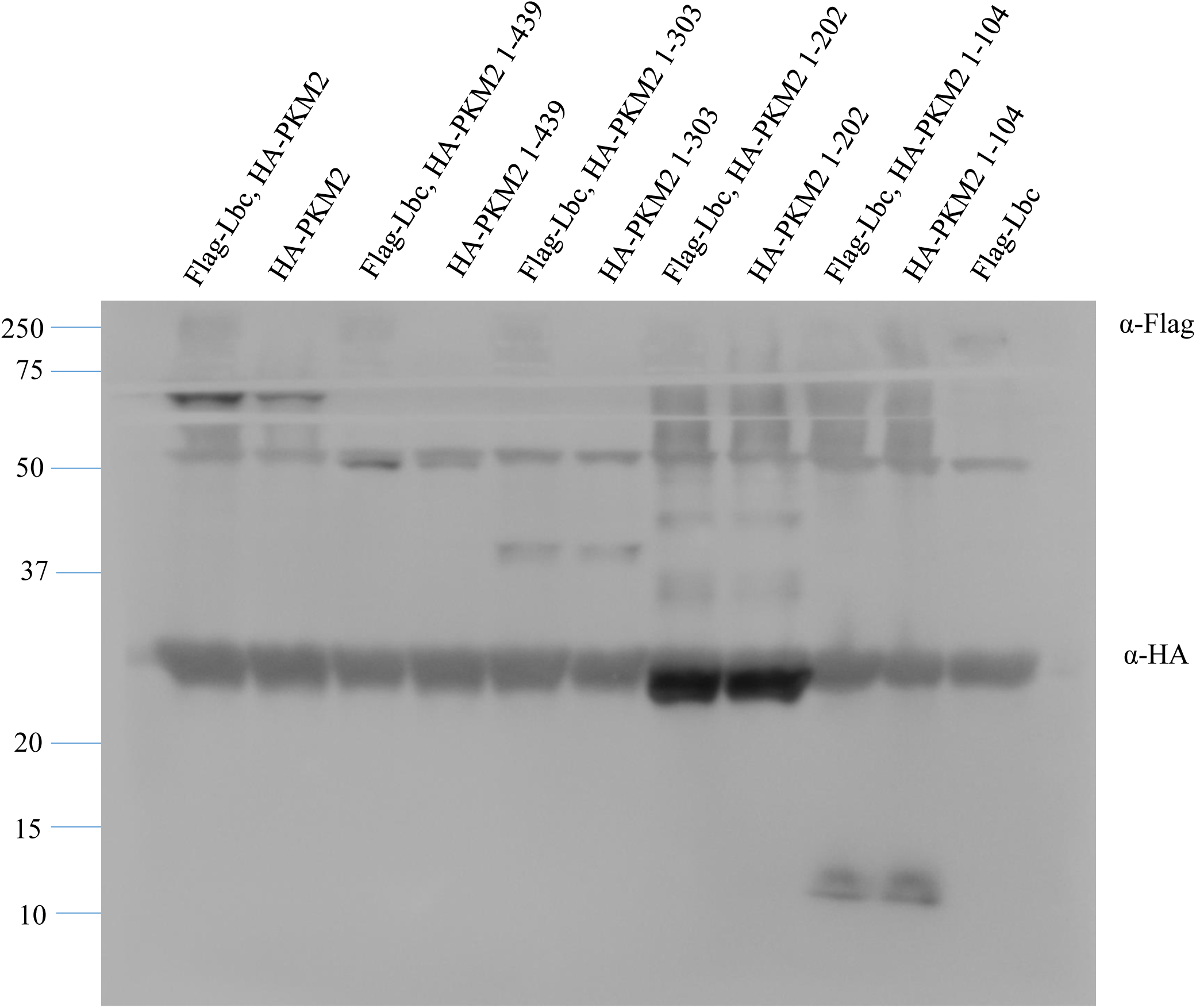
HA-PKM2 304-439 binds Flag-AKAP-Lbc. Anti Flag pull down to examine the binding domain in PKM2 to AKAP-Lbc.

### Characterization the binding domains of both proteins

To identify the binding domains of the proteins responsible for binding, we truncated both proteins based on their function domains (data not shown), and tested binding by pull down experiments. First, we expressed both Flag-AKAP-Lbc and HA-PKM2 fragments in HEK 293 cells, and then tried anti-Flag IP, then compared the amount of HA-PKM2 fragments with the loading control equal to 10% of that used for IP. We observed that anti-Flag beads pulled down much higher level of full length HA-PKM2 and HA-PKM2 1-439 compared to their corresponding loading controls but not for other fragments such as HA-PKM2 1-303, suggesting that PKM2 304-439 binds AKAP-Lbc (Figure 4A). Furthermore, we deployed anti-GST IP to explore the sequence of AKAP-Lbc which binds His-PKM2. In the assay, both GST-AKAP-Lbc fragments and His-PKM2 from purified after expression in E. coli, and we found that fragments 5 and 6, corresponding to AKAP-Lbc 1923-2366 and AKAP-Lbc 2337-2817, pulled down His-PKM2 (Figure 4B), suggesting the C-terminus of AKAP-Lbc binds PKM2 directly.

**Figure 4.**
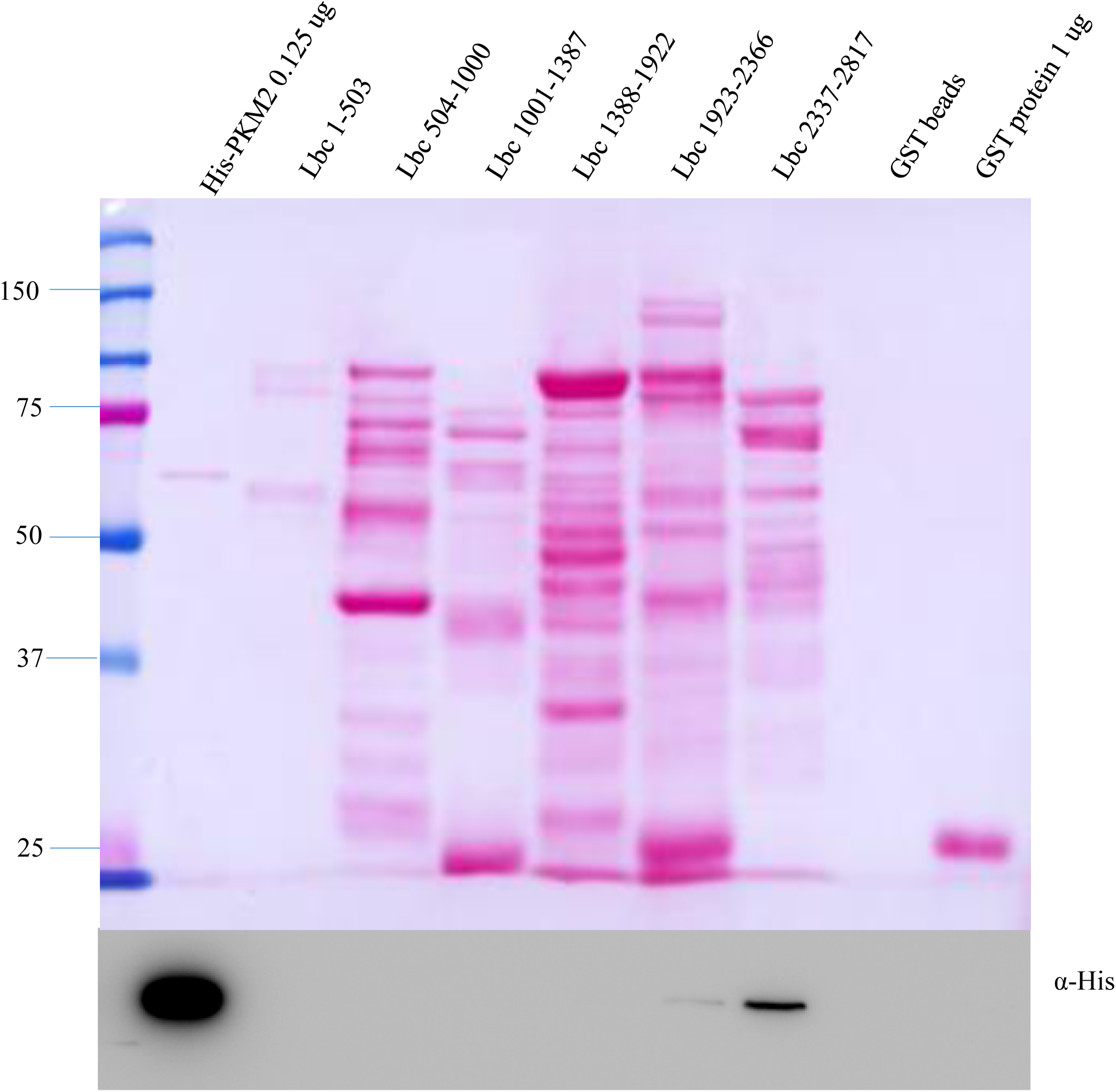
Identification of the interaction sequence of AKAP-Lbc to PKM2. Anti-Flag pull down with expression of both GST-AKAP fragments and His-PKM2. The upper panel shows the purified His-PKM2, GST-AKAP fragments and the GST beads and GST protein used for co-IP, while the lower ones exhibits the anti-His blot after pull down.

### Disruption of the binding between AKAP-Lbc and PKM2 inhibits cell growth

After determining the binding sequences of AKAP-Lbc and PKM2, we further asked whether we can interfere with binding by expressing fragments of both proteins. To our surprise, we did not observe that PKM2 fragments could significantly reduce the binding between these two proteins (data not shown). We next examined the effect of AKAP-Lbc fragments. We expressed Flag-AKAP-Lbc, HA-PKM2 and Myc-AKAP-Lbc fragments 1, 5 or 6 in HEK 293 cells, then after expression, the extracts were used for anti-Flag IP. Consistent with that AKAP-Lbc fragments bind PKM2 directly as shown in Figure 4B, we observed that fragments 5 and 6 but not 1 disrupted the binding between Flag-AKAP-Lbc and HA-PKM2 (Figure 5A); fragment 6 played more powerful role to block the binding, strongly implying that AKAP-Lbc fragment 6 may also influence the binding function of AKAP-Lbc and PKM2. To explore the possibility, we compared the effect of expressing Myc-AKAP-Lbc fragments 1 and 6, and found that both fragments inhibited cell growth while fragment 6 reduced cell proliferation more than fragment 1 (Figure 5B). Taken together, disruption of the binding between AKA-Lbc and PKM2 reduces cell growth.

**Figure 5.**
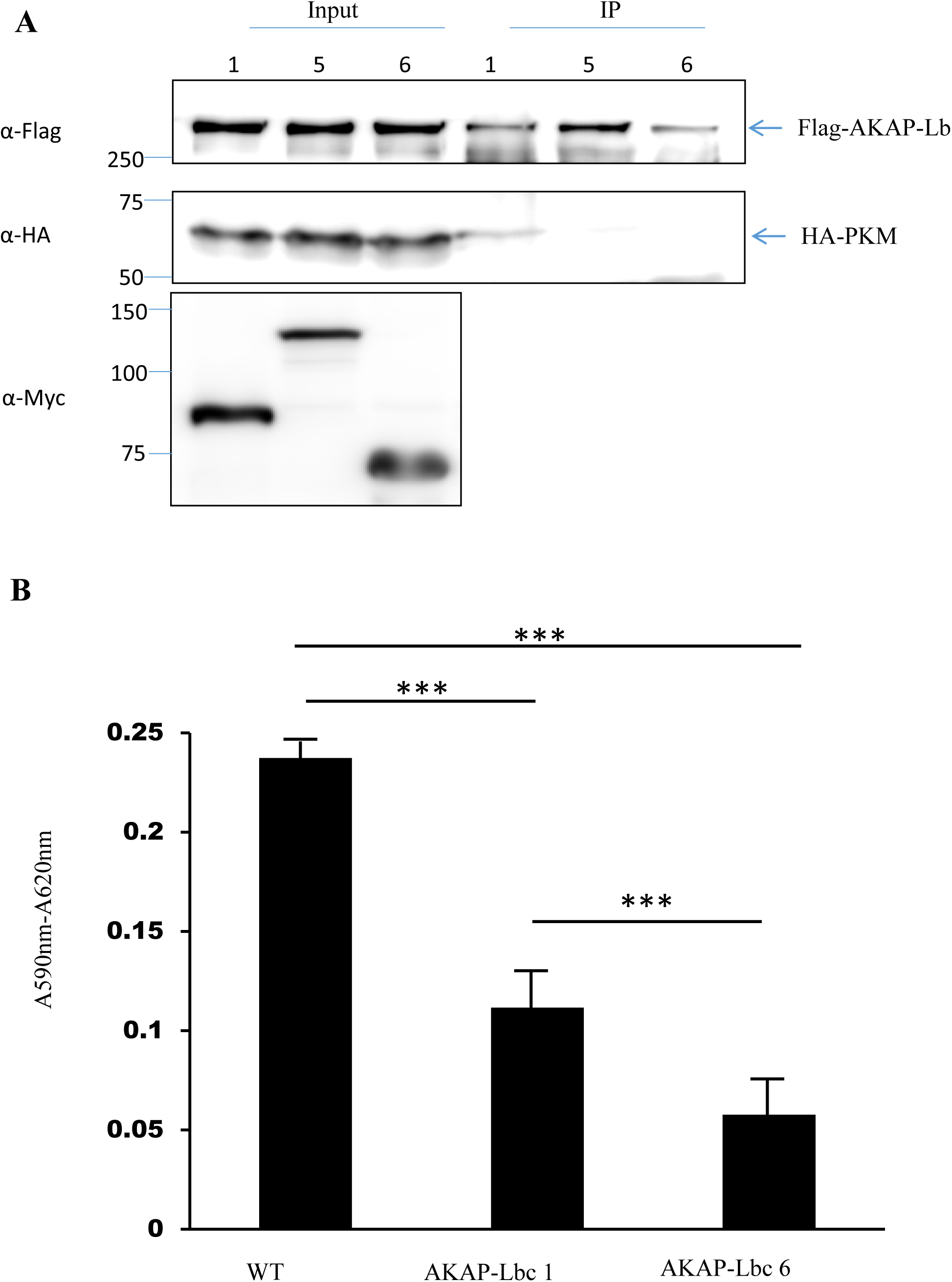
AKAP-Lbc fragments disrupt the binding between AKAP-Lbc and PKM2. Expression of Flag-AKAP-Lbc fragments 5 and 6 reduces the binding between AKAP-Lbc and PKM2. B) AKAP-Lbc fragment 6 inhibits cell proliferation much more than the fragment 1 compared with the full length of AKAP-Lbc.

## Discussion

In this work, we report that AKAP-Lbc and PKM2 bind directly, and the binding is important for cell growth.

Lbc was first characterized early in leukemias (Toksoz and Williams 1994), with a potential role in cancer biology as a transforming factor. Later, a longer variant called AKAP-Lbc was identified (Diviani, Soderling et al. 2001), and its roles in stress fiber formation and cardiac hypertrophy were discovered (Diviani, Soderling et al. 2001, Carnegie, Soughayer et al. 2008), while its potential function in cancer biology remains unclear. According to the yeast two-hybrid done by Alliance for Cell Signaling, AKAP-Lbc may interact with PKM2. Working as a scaffold protein, AKAP-Lbc provides a binding and interaction platform for other proteins or complexes to play their specific role in specified space (Diviani, Soderling et al. 2001), therefore it is attractive to propose PKM2 may bind AKAP-Lbc to regulate cell growth or proliferation or migration in cancer cells. To check the possibility, it is necessary to confirm that they really bind each other, and fortunately we observed the interaction.

To explore the biological function of the binding between proteins, it is practical to disrupt the interaction between proteins with polypeptides (Calejo and Tasken 2015). Normally, the specificity and efficiency of the blocking peptides decide whether we can design good ones to reduce the interaction between proteins. To improve the specificity of the peptides, it is primary to find the binding domains between proteins. We tried the PKM2 fragments to disrupt the binding between AKAP-Lbc and PKM2 but failed (data not shown), and we shifted to AKAP-Lbc fragments and found both fragments 5 and 6 interrupted the binding between the two proteins, while the fragment 6 worked more efficiently than 5 (Figure 5A), and we chose the fragments 1 and 6 to express in the HEK 293 cells to determine the biological function of binding between the two proteins. Fortunately, we found that both fragments can reduce cell proliferation, while fragment 6 reduces the proliferation much more (Figure 5B). All the results suggest that AKAP-Lbc plays an important role in cell proliferation by interacting with PKM2. Also, this work expands our understanding how PKM2 regulates cell proliferation, including its distribution mechanism and its new binding partner. More importantly, this work provides us a new strategy to lower cell proliferation, disruption of the binding between AKAP-Lbc and PKM2.

To further our insight of the binding between these two important proteins in cancer biology, it should be intriguing to explore the importance of the binding in other cancers, as well as other unique characters of cancer cells, such as migration, drug resistance, and so on. By that way, more potential roles of the binding might be uncovered, and the polypeptides blocking the binding might be optimized to treat some cancers.

## Acknowledgments

We are grateful to the support from American Heart Association Scientist Development Grant 11SDG5230003 and National Center for Advancing Translational Science-University of Illinois at Chicago Center for Clinical and Translational Sciences Grant UL1TR000050.

